# Quantifying turgor pressure in budding and fission yeasts based upon osmotic properties

**DOI:** 10.1101/2023.06.07.544129

**Authors:** Joël Lemière, Fred Chang

## Abstract

Walled cells, such as plants, fungi, and bacteria cells, possess a high internal hydrostatic pressure, termed turgor pressure, that drives volume growth and contributes to cell shape determination. Rigorous measurement of turgor pressure, however, remains challenging, and reliable quantitative measurements, even in budding yeast are still lacking. Here, we present a simple and robust experimental approach to access turgor pressure in yeasts based upon the determination of isotonic concentration using protoplasts as osmometers. We propose three methods to identify the isotonic condition – 3D cell volume, cytoplasmic fluorophore intensity, and mobility of a cytGEMs nano-rheology probe – that all yield consistent values. Our results provide turgor pressure estimates of 1.0 ± 0.1 MPa for *S. pombe*, 0.49 ± 0.01 MPa for *S. japonicus*, 0.5 ± 0.1 MPa for *S. cerevisiae W303a* and 0.31 ± 0.03 MPa for *S. cerevisiae BY4741*. Large differences in turgor pressure and nano-rheology measurements between the *S. cerevisiae* strains demonstrate how fundamental biophysical parameters can vary even among wildtype strains of the same species. These side-by-side measurements of turgor pressure in multiple yeast species provide critical values for quantitative studies on cellular mechanics and comparative evolution.

## Introduction

Turgor pressure is a primary determinant of mechanical properties of walled cells of various biological kingdoms including plants, fungi, bacteria, and protists (Shabala *et al*., 2009; Beauzamy *et al*., 2014). Turgor pressures also contribute to mechanical properties and cell shape determination in animal cell systems (Jones *et al*., 2021; Vian *et al*., 2022). Turgor pressure (also known as hydrostatic pressure) is the internal pressure relative to the outside environment of the cell; this internal pressure arises largely from the concentration of osmotic solutes such as ions, amino acids, and small metabolites. It is generated by an inward flow of water from a low solute concentration media into the cell that contains a higher solute concentration. This osmotic swelling pushes the plasma membrane against the cell wall. Consequently, the cell wall mechanically resists this turgor pressure (Abenza *et al*., 2015). Hence, there is an essential relationship between turgor pressure and cell wall mechanics.

Turgor pressure provides the force to drive the volume expansion for cell growth and contributes to cell shape determination and cytokinesis (Proctor *et al*., 2012; Chang and Huang, 2014; Atilgan *et al*., 2015). Walled cells harness turgor pressure to perform specific function, for example, some pathogenic fungi build appressoria that generate remarkably high turgor pressures that allows them to pierce and invade plant tissues or insects (Emmett and Parbery, 1975; Thilini Chethana *et al*., 2021). High turgor in fungi may also help maintain cell shape and integrity in compressive or arid environments in their natural habitat such as biofilms, soil, or decaying vegetative material (Mishra *et al*., 2022).

The internal pressure of the cell is highly relevant to membrane-based processes at the plasma membrane For example, internal pressure has been shown to impact the dynamics of endocytosis and cytokinesis, in yeasts as well as in animal cells (Aghamohammadzadeh and Ayscough, 2009; Proctor *et al*., 2012; Basu *et al*., 2013; Wagner and Glotzer, 2016; Li *et al*., 2017; Lemière *et al*., 2021; Wang *et al*., 2021). However, the precise contribution of hydrostatic pressure and how it is countered by actin-dependent forces remain poorly defined. Accurate measurements of turgor or hydrostatic pressure are critical to understand the mechanics of these plasma membrane processes at a quantitative level.

There is currently a critical need for reasonable estimates of turgor for quantitative modelling of the mechanics of plasma membrane dynamics and actin structures at the cell surface. Mechanical-based models of endocytosis and cytokinesis all depend on turgor pressure as a key parameter. For instance, reliable estimates of turgor are required to estimate the force production of actin filaments at the endocytic pit for endocytosis (Dmitrieff and Nédélec, 2015; Nickaeen *et al*., 2022).

Experimental measurement of turgor pressures in various cell types however has been highly challenging. There is no singular standard method; a variety of approaches for measuring turgor have been devised in different contexts and cell types and each has its caveats and challenges (Beauzamy *et al*., 2014). In the literature, various approaches could yield a large range of values (over one magnitude) even in the same cell type. One gold standard technique used in very large plant cells is to insert an oil-filled capillary into the cell as a direct pressure gauge (Tomos and Leigh, 1999). However, most cell types are not large enough or amenable to this sort of physical manipulation. Another approach involves using AFM-based cantilevers to press on the surface of cells and measure its resistance to displacement (Boulbitch, 1998; Beauzamy *et al*., 2015a; Tsugawa *et al*., 2022). One drawback of these indentation approaches is that analyses are needed to deconvolve the mechanical properties of the cell wall from the internal turgor pressure. Another class of approaches rely on measuring how cell volume changes in responses to osmotic shifts in the media (Atilgan *et al*., 2015). These rely on accurate 3D measurements of cell volume, and requires characterization of the intrinsic mechanical properties of the cell wall and cell shape beforehand (Atilgan *et al*., 2015).

Bacteria, fungi, plant cells, and zebrafish embryo measurements show that turgor pressure can approach megapascal values (Jiang and Sun, 2010; Beauzamy *et al*., 2015b; Goldenbogen *et al*., 2016; Altenburg *et al*., 2019; Vian *et al*., 2022); with reported values in cultured mammalian cells being ten times lower. In fission yeast *S. pombe*, Minc et al. (Minc *et al*., 2009) used cell buckling, and Atilgan et al. (Atilgan *et al*., 2015) used osmotic responses and modeling to arrive at consistent estimates in the range of 1.0-1.5MPa (or ∼10-15 atm or ∼145-217 Psi). These studies suggest that the fission yeast cell has an internal pressure similar to a racing bike tire, with the cell wall having a stiffness of hard rubber of ∼50 MPa (Atilgan *et al*., 2015). The published values for turgor pressure in *S. cerevisiae* however have been less consistent, with estimates ranging from 0.05 to 2.9 MPa (Arnold and Lacy, 1977; Martinez de Marañon *et al*., 1996; Meikle *et al*., 2009; Goldenbogen *et al*., 2016). Recent modeling of the endocytic pit derived from properties and organization of actin filaments and plasma membrane deformation back calculated a turgor pressure of 0.35 MPa (Nickaeen *et al*., 2022). Thus, accurate measurements of turgor pressure values in budding yeast are especially lacking.

Here, we introduce an approach for measuring turgor pressure based upon osmotic responses in protoplasts (yeast cells with their cell wall enzymatically removed). This method is based upon our previous findings that fission yeast protoplasts act as ideal osmometers; their osmotic potential is equal to the osmotic potential in the media outside of the cell (Lemière *et al*., 2022). The general premise is that by tuning the outside osmotic environment we adjust the protoplasts internal osmotic potential. We then determine the external osmolarity required for protoplasts to match the internal osmotic potential of intact cells. The outside osmolarity at this isotonic point provides an estimate of turgor pressure (Figure 1). Here, we developed three different ways to quantitatively pinpoint this isotonic state based on cell volume, cellular concentration, and cytoplasmic rheology. The three orthogonal approaches are advantageous in that they do not rely on prior knowledge of cell shape or cell wall mechanical properties. We derive consistent turgor measurements for *S. pombe*, *S. cerevisiae*, and *S. japonicus*, an emerging fission yeast model cell which is ten-fold larger in volume than *S. pombe*. One surprising finding is that two wildtype *S. cerevisiae* strains – W303 and BY4741, which represent the major strain backgrounds used in the field – differ in turgor pressure by almost two-fold, demonstrating how even "wildtype" cells of the same species can have substantial variability in their mechanical and osmotic properties.

**Figure 1.**
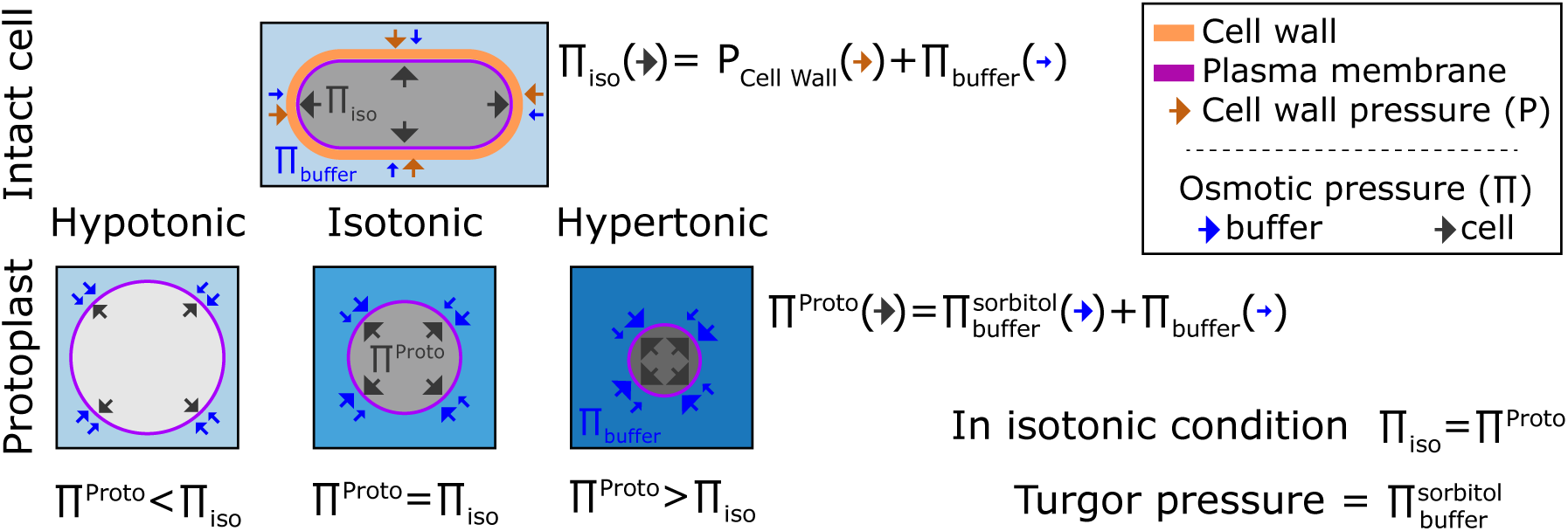
Representation of the turgor pressure measured with osmotic treatment. For intact cells in standard media, cells are in an isotonic condition in which pressures from the elastic cell wall (P_#(**,-**_) plus the osmotic buffer of the medium (∏_%&’’()_) counterbalance the internal pressure (∏_./0_). For protoplasts, the osmolarity of the external media causes the protoplast to increase or decrease in volume so that the internal osmotic pressure counterbalances pressure from external media. At isotonic condition, the internal pressure of the protoplast matches the internal pressure of the intact cell ∏^1)020^ = ∏_./0_. This isotonic pressure point is achieved when volume of the protoplast matches the volume of the intact cell. Thus, the turgor pressure of the intact cell can be estimated from sorbitol concentration in the external media needed to achieve this isotonic point in the protoplast.

## Results

### Protoplasts can be used as osmometers to gauge internal osmotic pressures

The isotonic condition is defined at the point where the water potential of a cell (Ψ_*C*_) is equal to the medium water potential (Ψ_*M*_) at equilibrium. In first order, there is a direct relation between the cell water potential, the internal osmotic potential (Π_*C*_), and its turgor pressure (P) such that Ψ_*C*_ = P − Π_*C*_. This relation is usually simpler for the medium where only the concentration of solutes is considered such that Ψ_*M*_ = −Π_*buffer*_. From these two equations, when cells are at equilibrium with the surrounding medium it comes a simple relation between the turgor pressure (P) and the difference of osmotic potential between the medium (Π_*buffer*_) and the cell (Π_*C*_): P = Π_*C*_ − Π_*buffer*_. The turgor pressure can then be seen as the mechanical resistance of the cell wall to expand due to the internal pressure inside a yeast (Figure 1).

Cells swell in hypotonic condition and shrink in hypertonic condition. From a mathematical point of view, this behavior comes from the fact that cells adjust their water potential to be at equilibrium with the surrounding medium. Because the membrane permeability is generally higher to water than to solutes cells adjust their water potential by exchanging water rather than solute (Lang *et al*., 1998; Milo and Phillips, 2015). Hence, under osmotic shocks at short time scales (< minute), the total amount of solute can be considered constant (Yang and Hinner, 2015).

A particular case emerges when cells do not bear any turgor pressure (P=0), such that the osmotic potential of the buffer and inside the cell are equal (Π_*C*_ = Π_%&’’()_). In this case, cells behave like ideal osmometers, whose volume varies with external osmotic pressure (Figure 1). The volume of an ideal osmometer shows a linear relationship with the inverse of the osmolytes concentration in solution, following the Boyle Van’t Hoff relation: V ∝ ^1^-(Nobel, 1969). Indeed, our recent study shows that *S. pombe* protoplasts exhibit %&’’() such properties of an ideal osmometer (Lemière *et al*., 2022).

We tested whether protoplasts from different yeast species all act as ideal osmometers. Protoplasts were generated by treating live yeast cells with cell wall digestive enzymes (see Methods). Protoplasts were distinguished from cells with an intact cell wall by their spherical cell shape and sensitivity to lysis at low osmotic conditions. After digestion of the cell wall, protoplasts were shifted into rich growth media with the indicated concentrations of sorbitol and then analyzed. To measure cell volume, we imaged cells expressing a fluorescent plasma membrane protein marker (mCherry-Psy1 for *S. pombe*, INA1-GFP for both *S. cerevisiae* strains, and Mtl2-mCherry for *S. japonicus* (Figure 2A)) and determined their cellular volumes using a semi-automated 3D-segmentation tool (Machado *et al*., 2019). When sorbitol was added to the growing medium, protoplasts decreased in volume. Inversely, reducing the amount of sorbitol swelled protoplasts to larger volumes. For each yeast strain, we plotted the volume of a population of protoplasts as a function of the inverse of the concentration in the medium (Figure 2B, bottom panel). The Boyle Van’t Hoff plots for protoplasts showed a linear behavior for all the yeast strains tested, indicating that protoplasts in the different strains all behaved as ideal osmometers and do not bear detectable turgor pressure.

**Figure 2.**
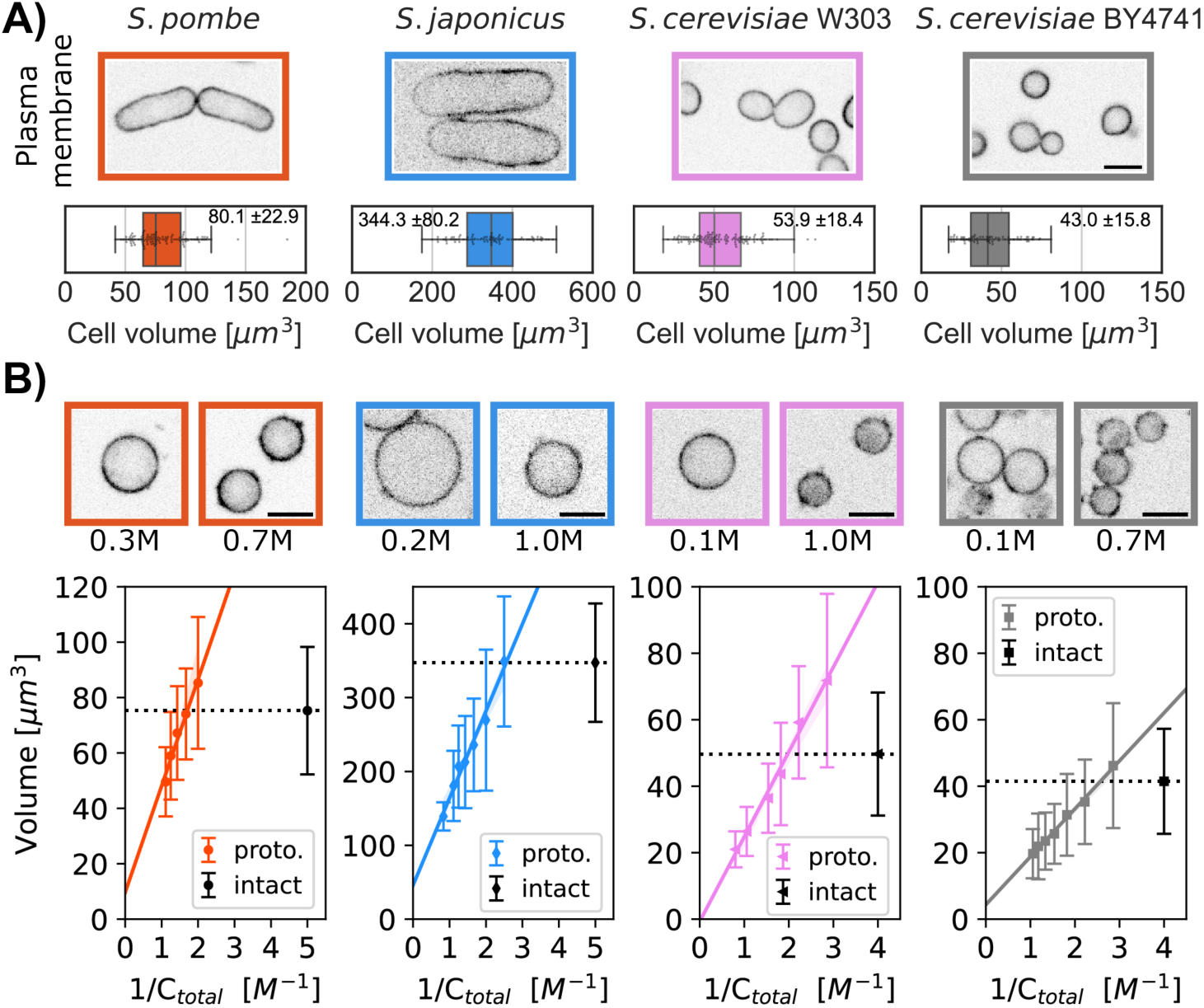
Comparison of the volumes of intact cells and protoplasts to quantify turgor pressure. A) Single confocal plane images of four yeast strains of the indicated species or background, each expressing a fluorescent plasma membrane marker. Graphs show the distribution of cellular volumes of normal asynchronous populations in growth medium (mean ± SD indicated in the box plot, N>91 cells). B) Single confocal plane images of protoplasts expressing a fluorescent plasma membrane marker in indicated sorbitol concentrations. Boyle Van’t Hoff plots show the relationships between protoplast volumes and osmolarity of the media (C_total_ = C_sorbitol_ + C_growth media_). The linear relationships indicate that the protoplasts all act as ideal osmometers. Dashed black lines on the plots represent the median cellular volume of a population of intact cells in growth medium without sorbitol. The colored lines represent the fit with the 95% confidence interval (A). The sorbitol concentration at which the dashed black line intersects with the protoplast volume (fitted colored line) is used to calculate turgor pressure values. Median ± SD, N>450 cells per strain, at least 2 replicates per point. Scale bar 5 µm.

We used protoplasts as ideal osmometers to determine the internal osmotic environment of intact cells. At the isotonic osmolarity, the volume of the protoplast matches the volume of the intact cell. Thus, to access turgor pressure, we measured the concentration of extracellular sorbitol needed to reach this isotonic state in the protoplast. In the following sections, we tested three approaches to determine this isotonic state.

### Measurement of turgor pressure with cell volume measurements

First, we directly measured the volume of cells (as described above, see Methods) to determine the isotonic volume condition. We treated protoplasts with a range of sorbitol concentrations and measured their volume distributions. We then compared these volume distributions to the volume distributions of intact cells grown in media without sorbitol (Figure 2B, top panel). Using the Boyle Van’t Hoff linear fit for protoplasts volume, we calculated the sorbitol concentration at which the mean volume of the protoplasts equaled the mean volume of intact cells (Figure 2B) (Table 1). We used the Van’t Hoff relation between the concentration of solute and the osmotic pressure ΙΙ_./0_ = ΙΙ_%&’’()_ = CRT, where RT is the product of the gas constant and the temperature (K) to access the turgor pressure. This method based upon 3D volume measurement yielded turgor pressures of 0.97±0.1 MPa for *S. pombe*, 0.48±0.06 MPa for *S. japonicus*, 0.64±0.07 MPa for *S. cerevisiae W303a* and 0.34±0.03 MPa for *S. cerevisiae BY4741* (Table 2).

**Table 1:** Sorbitol concentration (mol/L) at which protoplasts reach isotonic state compared to intact cells for the three different experimental approaches.

| **Strains**<br>**Methods** | ***S. pombe***<br>**(mol/L)** | ***S. japonicus***<br>**(mol/L)** | ***S. cerevisiae***<br>**W303 (mol/L)** | ***S. cerevisiae***<br>**BY4741 (mol/L)** |
| --- | --- | --- | --- | --- |
| **Cellular volumes** | 0.39± 0.4 | 0.19± 0.02 | 0.25± 0.03 | 0.13± 0.02 |
| **Cytoplasmic concentration** | 0.37± 0.3 | 0.21 0.03 | 0.16± 0.01 | 0.10± 0.01 |
| **Rheology** | 0.43± 0.02 | n/a | 0.20± 0.01 | 0.13± 0.01 |
| **Average values of these three methods** | 0.40± 0.02 | 0.20± 0.01 | 0.20± 0.04 | 0.12± 0.01 |

Summary of the sorbitol concentration added to the media for protoplasts to match the internal osmotic potential of intact cells for the three methods (see Methods). Last row gives the average values from the three methods (Mean ± SD).

**Table 2:** Results of turgor pressure measurements in MPa for the three different experimental approaches in this study.

| **Strains**<br>**Methods** | ***S. pombe***<br>**(MPa)** | ***S. japonicus***<br>**(MPa)** | ***S. cerevisiae***<br>**W303 (MPa)** | ***S. cerevisiae***<br>**BY4741 (MPa)** |
| --- | --- | --- | --- | --- |
| **Cellular volumes** | 0.97± 0.1 | 0.48± 0.06 | 0.64± 0.07 | 0.34± 0.03 |
| **Cytoplasmic concentration** | 0.95± 0.1 | 0.50± 0.06 | 0.43± 0.03 | 0.27± 0.02 |
| **Rheology** | 1.09± 0.09 | n/a | 0.51± 0.03 | 0.32± 0.02 |
| **Average values of these three methods** | 1.0± 0.1 | 0.49± 0.01 | 0.5± 0.1 | 0.31± 0.03 |

Summary of the turgor pressure computed for the three methods (see Methods). Last row gives the average values from the three methods (Mean ± SD).

### Measurement of turgor pressure using fluorescence intensity measurements

Second, we assessed cell volume indirectly by measuring fluorescence intensity of a cytoplasmic marker. One drawback of the volume measurement approach described above is that it is based upon distributions in an asynchronous population, and thus care must be taken not to under sample. The measurement of intracellular intensity has the advantage of an intensive property: it does not depend on the number of cells assayed.

A fluorescent cytoplasmic protein marker was expressed in each of the strains *S. pombe* (m-Crimson from Act1 promoter), *S. japonicus* (GFP from Adh1 promoter), and *S. cerevisiae* (td-Tomato from CAN promoter) from constructs integrated into the chromosome (see Methods) (Table 3). We measured the fluorescence intensity of the fluorophore in a small region of interest within each cell or protoplast in a representative portion of the cytoplasm (see Methods). We verified that the fluorescent intensities of these cytoplasmic proteins did not vary through the cell cycle (Figure 3A). In hypertonic conditions, the mean intensities of the cytoplasmic fluorophore increased as the protoplasts volume decreased, while intensities decreased in hypotonic conditions when the protoplasts swelled (Figure 3B top panel) (Lemière *et al*., 2022; Molines *et al*., 2022).

**Figure 3.**
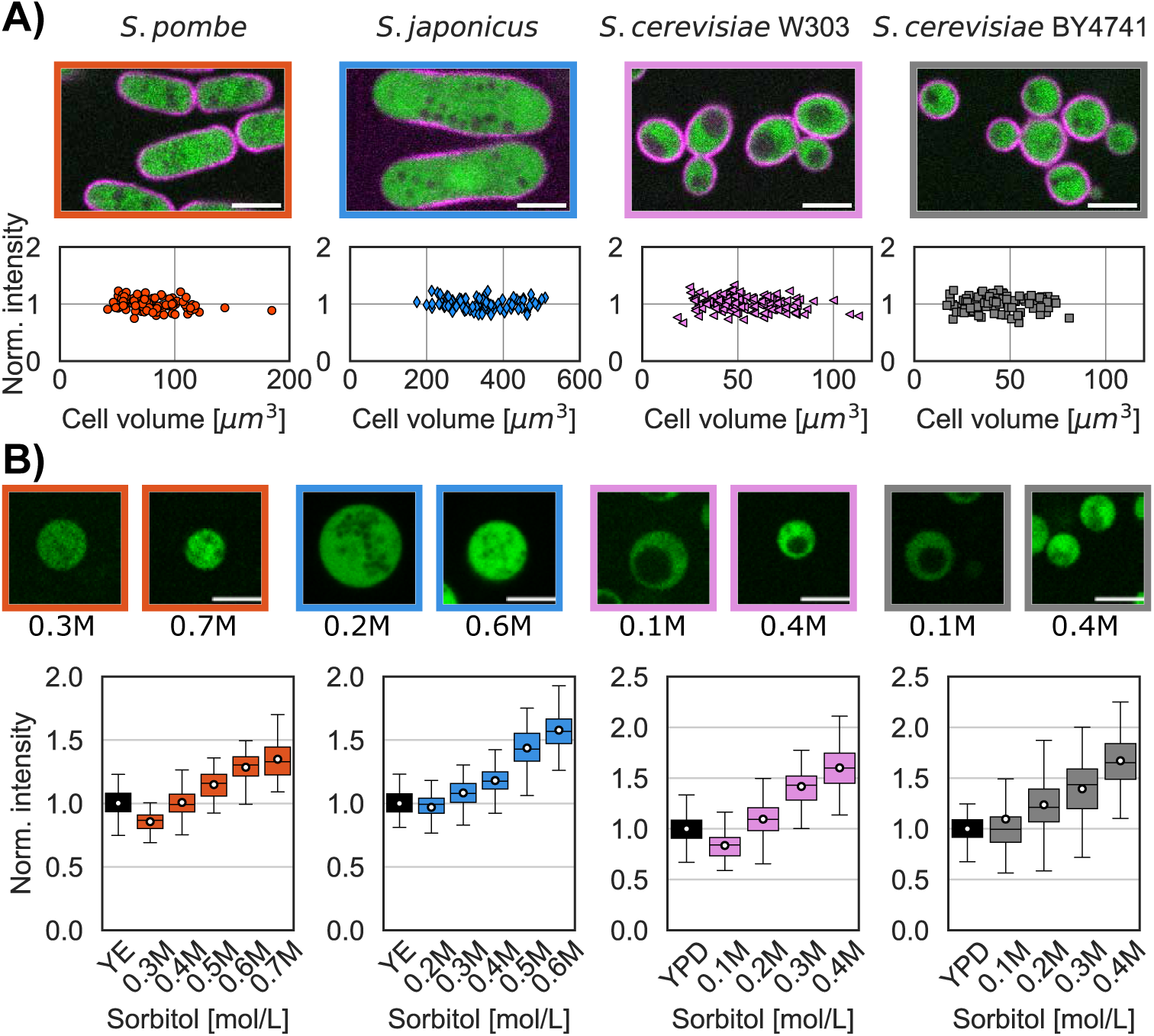
Measurement of turgor pressure with a cytoplasmic marker. A) Single confocal plane images of the 4 yeast strains expressing a plasma membrane marker (magenta) and a cytoplasmic marker (green). Graph shows the distribution of fluorescence intensity of the cytoplasmic marker in intact cells, showing that the concentration of these markers is maintained at a constant level through the cell cycle. B) Images of protoplasts expressing a cytoplasmic marker in the lowest and highest sorbitol concentrations tested for each strain. Graphs show the cytoplasmic intensity distribution of a population of intact cells (black whisker box) compared with the intensity in protoplasts in media supplemented with indicated concentrations of sorbitol (at least 2 replicates per condition, white dots mean values, N>400 cells per strains). The higher sorbitol concentration leads to increased fluorescent intensity in the cells due to higher concentration of the cytoplasmic marker. Scale bar 5 µm.

**Table 3:**
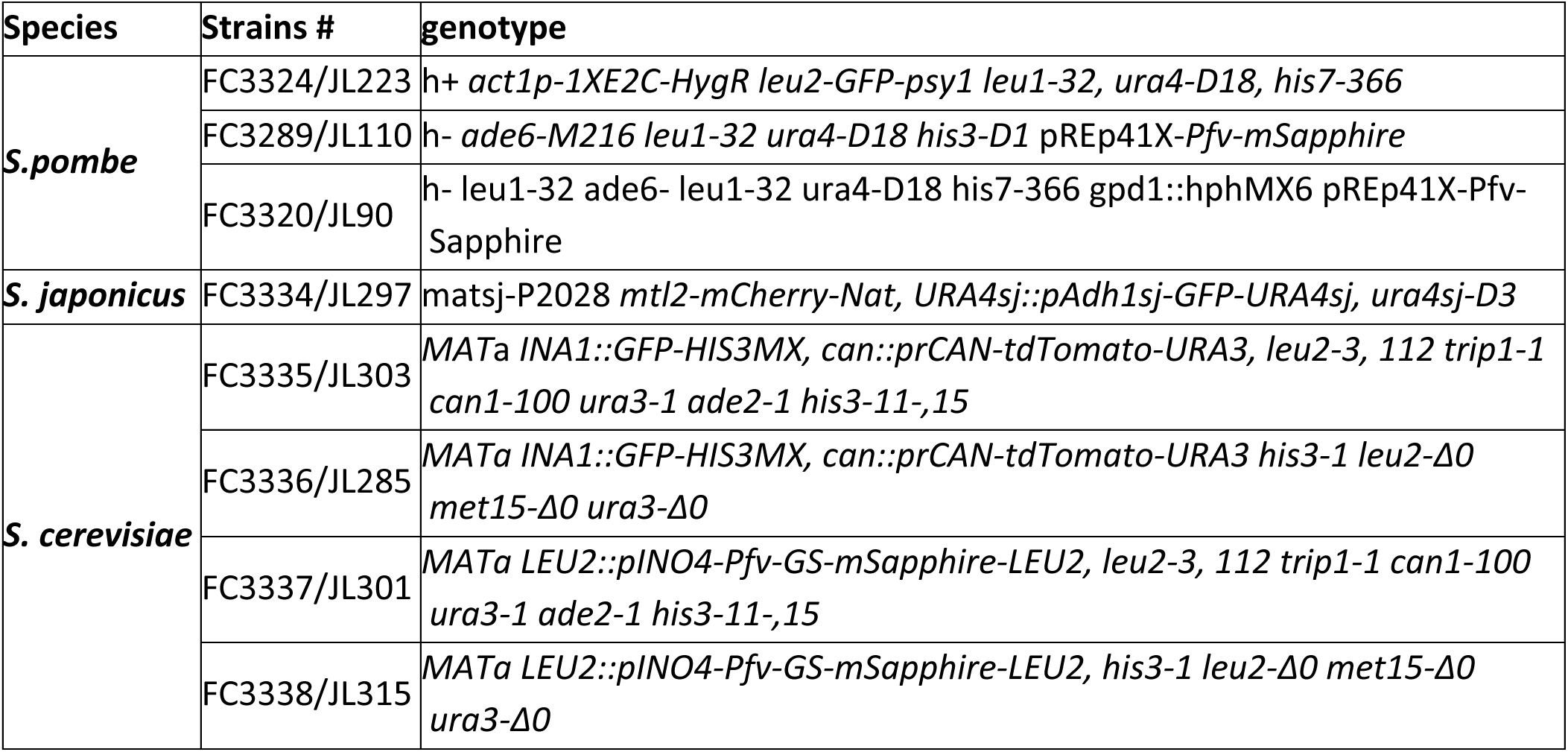
Strain list.

We monitored the fluorescence intensities in a population of protoplasts in their growth medium supplemented by various concentrations of sorbitol and compared these to intensities in intact cells. We calculated the sorbitol concentrations that lead the intensity in protoplast to match that in intact cells in the isotonic condition (Table 1). Using this method, we obtained turgor pressure values of 0.95±0.1 MPa for *S. pombe*, 0.50±0.06 MPa for *S. japonicus*, 0.43±0.03 MPa for *S. cerevisiae W303a* and 0.27±0.02 MPa for *S. cerevisiae BY4741* (Table 2).

### Measurement of turgor pressure using nano-rheology

The third method of determining the isotonic condition consisted of using nanorheology to measure diffusion within the cytoplasm (D_eff_). As probes to measure diffusion coefficients, we expressed 40-nm sized genetically-encoded multimeric cytoplasmic nanoparticles (cytGEMs) labelled with mSapphire-fluorescent protein (Delarue *et al*., 2018; Lemière *et al*., 2022; Molines *et al*., 2022). The mobility of cytGEMs in *S. pombe* and *S. cerevisiae* protoplasts followed a physical model of self-diffusive particles in a polymer solution (Figure 4B) (Phillies, 1988; Lemière *et al*., 2022), yielding a model that quantitatively relates D_eff_ with sorbitol concentration and cell volume during osmotic shifts experiments.

**Figure 4.**
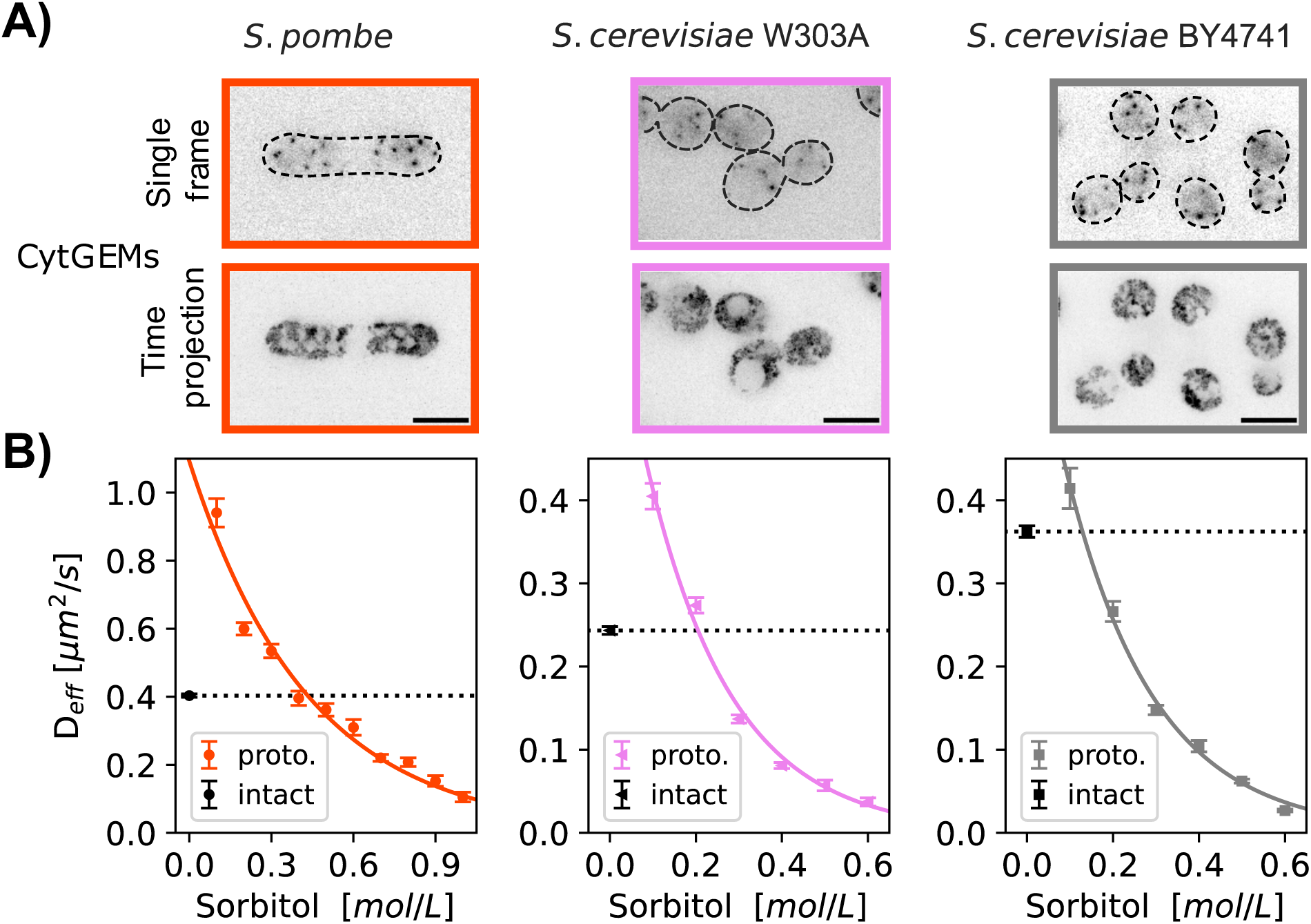
Measurement of turgor pressure using nano-rheology probes. A) Fluorescence images of cells expressing cytGEMs nanoparticles (inverted contrast). Top, single time point images with cell boundaries outlined (dashed lines); bottom, maximum time projections (500 time-frames over 5 sec). B) cytGEMs effective diffusion (D_eff_, Mean values ± SEM, N>5200 tracks per strain, at least 2 replicates per point) of intact cells in their standard growth media (black symbols and dashed lines) and of protoplasts in growth media supplemented with various sorbitol concentrations. cytGEMs D_eff_ values for protoplasts were fitted to a Phillies model (colored line), which was used to calculate the sorbitol concentration for which D^.=2->2^ = D^3)0203*-/2^, giving access to the turgor pressure values.

We expressed cytGEMs in the *S. pombe* and *S. cerevisiae* strains, but unfortunately, were not successful in expressing the protein in *S. japonicus*. We quantified cytGEMs mobility in each population of exponentially growing intact cells in their respective media at 30°C. For WT *S. pombe,* D^304%(^= 0.40 µm^2^/s, similar to values previously reported (Lemière *et al*., 2022; Molines *et al*., 2022). For *S. cerevisiae* strains, cytGEMs mobility for BY4741 was D^567879^ = 0.36 ± 0.01 µm^2^/s while for the W303 background D^,:;:<^ = (’’ (’’ 0.24 ± 0.01 µm^2^/s. These statistically significant differences (Mann-Whitney test, p<0.0001) suggest that diffusion is slower in W303 than in BY4741 and *S. pombe*.

We then determined the mobility of cytGEMs for protoplasts in media supplemented by various concentration of sorbitol. We found that the mobility of cytGEMs in *S. pombe* protoplasts and *S. cerevisiae* both fit a physical model that related D_eff_ to sorbitol concentration (Phillies, 1988; Lemière *et al*., 2022) (Figure 4B). Using this model, we calculated the sorbitol concentration needed to match cytGEMs mobility in the population of protoplasts with intact cells in the isotonic condition. Using this nano-rheology method, we calculated turgor pressure values:1.09±0.09 MPa for *S. pombe*, 0.51±0.03 MPa for *S. cerevisiae W303a* and 0.32±0.02 MPa for *S. cerevisiae BY4741*.

To test whether our turgor measurements may be altered by osmotic stress responses during the protoplasting procedure, we measured turgor in a fission yeast *gpd1Δ* mutant, which is defective in turgor adaptation in response to hyperosmotic shocks (Aiba *et al*., 1995; Lemière *et al*., 2022). We found that using cytGEMs in protoplasts, *gpd1Δ* mutant cells exhibited a similar turgor pressure as WT cells (1.05± 0.02 MPa in *gpd1Δ*, derived data from data in (Lemière *et al*., 2022)). Thus, at least in fission yeast, our turgor measurements were not affected by turgor adaptation.

Table 2 lists the collective results for the three experimental approaches in *S. pombe*, *S. japonicus, and S. cerevisiae*. *S. pombe* displayed the highest turgor pressure of the group, with the values consistent with recent measurements using other approaches (Minc *et al*., 2009; Atilgan *et al*., 2015). As each methods carried its own technical challenges, it was reassuring that the values were relatively consistent across the different methods without apparent systematic biases. Thus, these results establish these methods as robust approaches for the measurement of turgor pressure.

## Discussion

Here, we develop new methods for measurement of turgor pressure in yeast cells. This approach is based upon determining the isotonic environment that maintains the normal volume and cytoplasmic concentration in protoplasts compared to intact cells. Three methods for determining the isotonic protoplast volume – by measuring cell volume, cytoplasmic concentration, and nano-rheology – all yielded similar values, demonstrating the robustness of this approach. This study establishes turgor pressure values for *S. cerevisiae* (0.3-0.6 MPa) and *S. japonicus* (∼0.5 MPa) and *S. pombe* (∼1.0 MPa) (Table 2). These values represent a significant advance for the quantitative understanding of cell mechanics and have broad implications in determining forces in processes such as cell shape, cell size and growth, cytokinesis, and endocytosis. As these measurements were obtained together using the same methods, they provide a robust comparison between these species and strain background.

Our methods have advantages over other approaches such as osmotic shifts or cantilever indentation on walled cells (Routier-Kierzkowska *et al*., 2012; Atilgan *et al*., 2015) in that they do not depend on cell shape or detailed knowledge of cell wall-surface mechanical properties. Rather than trying to derive absolute values, our approach relies on matching values of the intact cell to a calibrated standard curve based on protoplasts. Our current methods yield average values for a population of cells but do not inform on cell-cell variability. Each of these three approaches to determine the iso-osmotic concentration of protoplasts compared to walled cells has its experimental advantages and challenges, depending on the expertise and local resources of the user. For 3D volume measurements, there are recognized challenges in segmentation for volume calculations and a need for large sample size to obtain distributions in asynchronous cells. Fluorescence intensity measurements depend on careful microscope calibrations and are sensitive to microscopy conditions (Methods). Measurements of cytGEMs mobility, which might currently be the most robust of these, require the introduction of cytGEMs into cells, a temperature controlled imaging system suitable for rapid acquisition (10-30 frames per second) and tracking analyses.

Our protocol utilizes protoplasts because they are ideal osmometers, in contrast to cells with intact cell walls which are not. However, the use of protoplasts comes with a potential caveat that during the procedure to generate them, cells may alter their internal osmolarity by synthesizing intracellular glycerol in response to stress. A number of observations suggest though that the internal osmolarity does not change during the protoplasting procedure. First, the turgor pressure values we obtain from fission yeast protoplasts are consistent with previous measurements using different approaches in intact cells (Minc *et al*., 2009; Atilgan *et al*., 2015). Second, fission yeast *gpd1* mutant protoplasts, which are defective in the synthesis of glycerol in response to osmotic stress, display similar turgor pressures and cytGEMs diffusion to WT cells using this method (Lemière *et al*., 2022), showing that measurements were not altered by turgor adaptation during the protoplasting procedure.

Our comparison between the two fission yeast species *S. pom*be and *S. japonicus* provides insights into how various mechanical parameters are altered to accommodate changes in cell sizes during evolution. The stress on the cell wall, for instance, is dependent on the radius of the cell (see Methods). *S. japonicus* has a similar rod-shape as *S. pombe* but is 2-fold larger in cell width and ∼10-fold larger in volume. The *S. japonicus* cell wall is approximately two times thicker than that in *S. pombe* (Davì *et al*., 2019). Experiments on measuring cellular dimensions upon cell lysis indicate that these species have similar cell wall elastic strain (Davì *et al*., 2019). Our measurements show that *S. japonicus* has two times less turgor pressure compared to *S. pombe* (Table 2). These values suggest that the cell wall in both fission yeasts support a similar tension of ∼1N/m (see Methods). These values allow us to derive a Young’s modulus (elasticity) of the lateral cell wall to be Y ∼ 50 MPa for *S. pombe* (Atilgan *et al*., 2015) and ∼ 25 MPa for *S. japonicus.* Thus, the ten-fold larger *S. japonicus* has similar cell wall tension as *S. pombe*, produced by half the turgor pressure that is supported by a thicker and more flexible cell wall.

One of the most unexpected findings was the approximately two-fold differences in turgor pressure and cytGEMs diffusion between two widely used *S. cerevisiae* strains BY4741 and W303. BY4741 is a derivative of the S228c strain that was used for the reference genome sequencing (Mortimer and Johnston, 1986; Brachmann *et al*., 1998). W303 is derived from multiple backgrounds and has substantial regions of the genome that originate from strains not related to S288c (Rothstein and Sherman, 1980b, 1980a; Rogowska-Wrzesinska *et al*., 2001). Comparison of the genome sequences of W303 and S228c strains show about 8000 nucleotide differences, predictive of amino acid changes in about 800 proteins (Ralser *et al*., 2012; Matheson *et al*., 2017). Numerous phenotypic differences between the strains have been noted, including cell size, salt tolerance, and maximal life span (Cohen and Engelberg, 2007; Zadrag-Tecza *et al*., 2009; Petrezselyova *et al*., 2010) (also see Figure 2A). The phenotypic differences in these two strains provides an opportunity to dissect molecular factors that regulate turgor pressure and cytoplasmic crowding. For instance, according to the Saccharomyces Genome Database, W303 carries an extra gene copy of ENA/ PMR2 (Cherry *et al*., 2012), which encodes plasma membrane ATPase pumps that regulates ion homeostasis which likely affects turgor pressure (Wieland *et al*., 1995). GEMs diffusion is thought to reflect the concentration of macromolecular crowding agents such as ribosomes (Delarue *et al*., 2018) rather than small molecules in the cytoplasm responsible for turgor pressure determination. Thus, W303 might have a more dilute cytoplasm with lower concentration of macromolecules. At present, it is unclear whether this significant difference in cytGEMs diffusion is somehow linked to the turgor pressure effect or is an independent effect.

Despite these physical differences, these yeast strains have both been selected to be robustly healthy and fast growing, at least in laboratory conditions. Although it has been long assumed that cells need to maintain optimal levels of turgor pressure and cytoplasmic properties (Minton, 1981), our findings suggest that there may not be a specific optimal level of turgor pressure and crowding, even within cells of the same species. Differences in turgor pressure and diffusion are however predicted to give rise to many physical and molecular differences driving biological processes such as endocytosis, cell growth, and cell size. Indeed, according to theory developed by Nickaeen et al. (Nickaeen *et al*., 2022), differences in turgor pressure predict that endocytosis may occur more quickly and require more actin in *S. cerevisiae* W303A than in BY4741, and similar differences in *S. pombe* compared to *S .japonicus.* Indeed, turgor pressure and rheological differences may begin to explain sources of variability between strains in the literature. The establishment of robust assays will stimulate further quantitative studies into the regulation of cell rheology and mechanics and their effects on cellular functions.

## Materials and Methods

### Yeast strains and media

Strains used in this study are listed in Supplemental Table 3. *S. pombe* strains are derived from WT 972 and *S. japonicus* strain is derived from NIG5091. Primers used to insert tagged genes are listed in Supplemental Table S2. For *S. pombe* and budding yeast, transformations were done with Lithium Acetate-based methods (Bähler *et al*., 1998; Longtine *et al*., 1998) (Table 4). *S. japonicus* was transformed using an electroporation method (Aoki and Niki, 2017). For expression of Cytoplasmic 40-nm GEMs in *S. pombe*, Pfv encapsulin-mSapphire was expressed from a thiamine regulated *nmt1*\* promoter on a multicopy plasmid pREP41X-Pfv-GS-mSapphire (Delarue *et al*., 2018; Lemière *et al*., 2022; Molines *et al*., 2022). For *S. cerevisiae* W303A background, cytGEMs integrative vector (#116930, pRS305-Leu2-PINO4-PfV-GS-mSapphire, Addgene) was inserted in the LEU2 locus after linearization with EcoRV restriction enzyme. As this strategy cannot apply to *S. cerevisiae* BY4741 strains as the LEU2 locus was completely deleted, we used a standard PCR-based method (Bähler *et al*., 1998) to insert pINO4::Pfv-GS-mSapphire-LEU2 into the region of the deleted LEU2 using primers matching the flanking region of the LEU2 deleted site including the restriction site added when this strain was designed (Brachmann *et al*., 1998) (see Table 4 for primers).

**Table 4:**
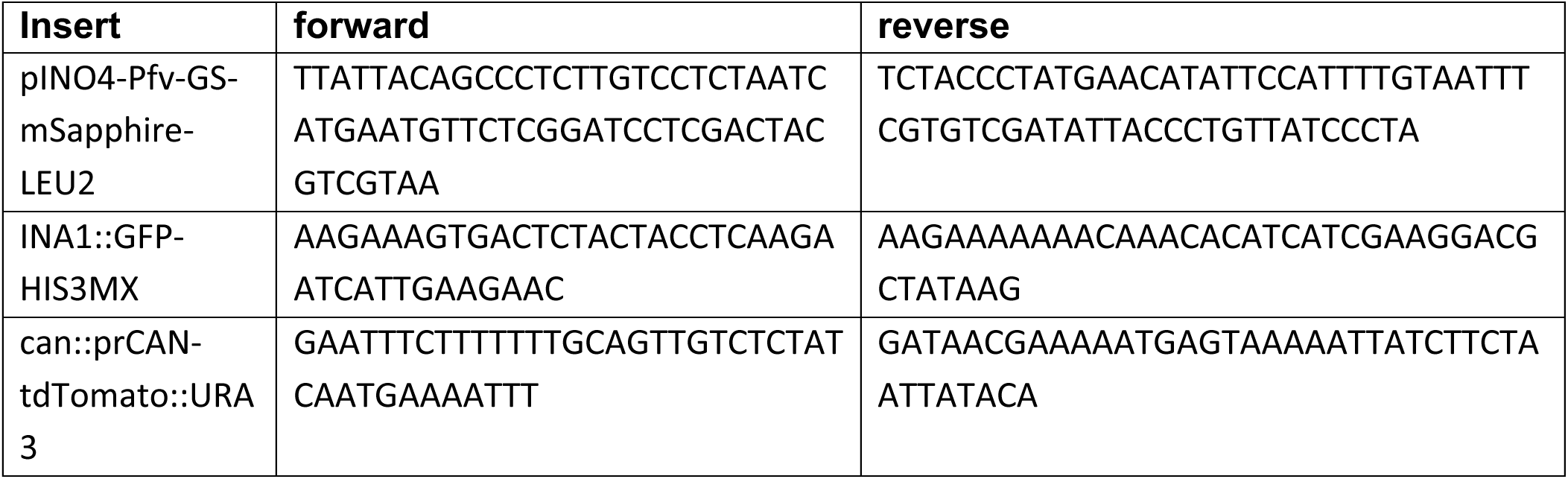
Primers.

*S. pombe* and *S. japonicus* cells were grown in YES 225 (#2011, Sunrise Science, San Diego CA, USA), *S. cerevisiae* cells were grown in YPD (10 g/L Bacto Yeast Extract, 20 g/L Bacto Peptone (#214010 and #0118-17-0, Becton Dickinson, Franklin Lakes NJ, USA, 20 g/L D-Glucose (#47263, Thermo Fisher Scientific, Waltham MA, USA)), all at 30°C in exponential phase for about 20 hr. For *S. pombe* strains carrying GEMs expression plasmids were grown in similar conditions in EMM3S – Edinburgh Minimum media (#4110-32, MP Biomedicals, Burlingame CA, USA) supplemented with 0.225 g/L of uracil, histidine, adenine, and 0.05 µg/mL of thiamine (#U0750, #H8000, #A9126, #T4625, Sigma-Aldrich, Saint-Louis MO, USA) to maintain the plasmid.

### Protoplast preparation

The protocol to produce fission yeast protoplasts is similar to the one described in (Flor-Parra *et al*., 2014; Lemière *et al*., 2022). *S. pombe* and *S. japonicus* cells were inoculated from fresh YES 225 agar plates (YES 225 + 20g/L Difco Agar (#281210, Fisher scientific)) into YES 225 or EMM3S liquid cultures and grown at 30 °C for about 20 h into exponential phase (OD600=0.2–0.3). Ten mL of cell culture was harvested by centrifugation 2 min at 400 rcf, washed two times with SCS buffer (20 mM sodium citrate, 20 mM citric acid, 1 M D-sorbitol, pH = 5.8), resuspended in 1 mL of SCS buffer with 0.1 g/ mL Lallzyme (#EL011-2240-15, Lallemand), and incubated with gentle inversion on a rotator for 10 min at 37 °C in the dark. The resulting protoplasts were gently washed once with 1 mL of SCS buffer using centrifugation for 2 min at 400 rcf. 900 μL of supernatant were removed, and the protoplasts in the pellet were gently resuspended in the remaining ∼100 μL of solution. This protoplasting procedure took about 15 min.

The S*. cerevisiae* protoplast protocol is similar to the one provided by MP Biomedicals white paper on Zymolyase. Cells were inoculated from fresh YPD agar plates (YPD +15g/L Bacto Agar (#214010, Fisher Scientific)) into liquid YPD and grown in exponential phase for 20 hours at 30°C. 15 mL of cells was harvested at 4000 rpm for 5 min and resuspended in 300 µL of TE Buffer (100 mM Tris pH 8.0, 100 mM EDTA). 400 µL of water and 3.5 µL of beta-mercaptoethanol were added, and cells were incubated at 30°C with gentle shaking for 15 min. Cells were washed once, centrifuged at 4000 rpm for 5 min. and resuspended in 800 µL of S buffer (1 M sorbitol, 10 mM PIPES). 8 µL of 100T Zymolyases from *Arthrobacter luteus* (#E1004, Zymo Research, Irvine CA, USA) was added, and cells were incubated at 30°C with gentle inversion on a rotator for 30 min. Protoplasts were harvested by centrifugation (5000 rpm, 5 min) and resuspended in S buffer. This protoplasting procedure took about 1 hour.

### Microscopy

Cells were imaged on a Ti-Eclipse inverted microscope (Nikon Instruments, Melville, NY, U.S.A) with a spinning-disk confocal system (Yokogawa CSU-10) that includes 488 nm and 541 nm laser illumination Borealis system and emission filters 525±25 nm and 600±25 nm respectively, a 60 X (NA: 1.4) objective, and an EM-CCD camera (Hamamatsu, C9100-13). These components were controlled with μManager v. 1.41 (Edelstein *et al*., 2010, 2014). Temperature was maintained by a black panel cage incubation system (#748–3040, OkoLab, Sewickley - PA, USA).

For imaging of GEMs, live cells were imaged using highly inclined laser beam illumination on a TIRF Diskovery system (Andor, Concord, MA 01742, USA) with a Ti-Eclipse inverted microscope stand (Nikon Instruments), 488 nm laser illumination, a 60 X TIRF oil objective (NA:1.49, oil DIC N2) (#MRD01691, Nikon), and a sCMOS camera (Zyla, Andor), controlled with μManager v. 1.41 (Edelstein *et al*., 2010, 2014). Temperature was maintained by a black panel cage incubation system at 30°C (#748-3040, OkoLab). 500 images were acquired at 100 fps with 10 ms exposure time with no interval time.

Cells were mounted in μ-Slide VI 0.4 channel slides (#80606, Ibidi – 6 channels slide, channel height 0.4 mm, length 17 mm, and width 3.8 mm, tissue culture treated and sterilized). For *S. pombe* and *S. japonicus* cells and protoplasts, the μ-Slide channel was pre-coated by incubation with 100 μg/mL of lectin from soybean (#L1395, Sigma) for at least 15 min at room temperature. For *S. cerevisiae* cells and protoplasts, μ-Slide channels were coated with 1 mg/mL of concanavalin A (#L7647, Sigma) until dried and could be stored at 4°C for days. Cells were introduced into the chamber already mounted under the microscope, incubated for 5 min to let them sediment and adhere to the chamber. The chamber was then washed three times, in less than a minute, to remove non-adhered cells. The media used for these washes and the final media contained the growth medium (YES 225 or YPD) with various amount of sorbitol. Then up to 5 different fields of view for each strain were acquired in < 1 min.

### 3D volume measurements

Cell volumes were measured by imaging cells expressing plasma membrane markers. For *S. pombe* we used mCherry-Psy1 (Kashiwazaki *et al*., 2011); for *S. japonicus* Mtl2-mCherry (Davì *et al*., 2018); *for S. cerevisiae* Ina1-GFP (Huh *et al*., 2003). Z stack images (0.5 μm z-slices) that spanned the entire cell volume were obtained using spinning disk confocal microscopy. The 3D volumes were segmented using an ImageJ 3D image segmentation tool LimeSeg (Schneider *et al*., 2012; Machado *et al*., 2019).

### Cytoplasmic marker intensity measurements

A fluorescent cytoplasmic protein marker was expressed in *S. pombe* (m-Crimson expressed from Act1 promoter), *S. japonicus* (GFP expressed from Adh1 promoter), and *S. cerevisiae* (td-Tomato expressed from CAN promoter). Cells were imaged with a spinning-disk confocal system only once to avoid photobleaching. Multiple fields of view were acquired per condition. The distance between the air-glass interface and the objective was locked during the entire set of experiments for a given μ-Slide. The cytoplasmic intensity was measured for each cells using the mean fluorescence intensity of a ROI manually selected at a given Z-position in the cells avoiding the vacuoles and corrected by the mean fluorescence of the background. To calibrate and normalize the intensity of our measurement throughout days and samples, at the beginning of each set of experiments for a given strain, a population of intact cells was imaged and analyzed in the exact same conditions (same laser intensity, exposure time and Z-position). Intensities of that intact cell population, for a given strain, was then used to normalize the set of experiments on protoplasts of the same strain for a given μ-Slide channel (the 5 other channels) such that one ibidi µ-Slide was used per strain and per replicates. This calibration was done for each µ-Slide and strains imaged.

### cytGEMs analyses methods

cytGEMs were tracked with the ImageJ Particle Tracker 2D-3D tracking algorithm from MosaicSuite (Sbalzarini and Koumoutsakos, 2005) with the following parameters: run("Particle Tracker 2D/3D", "radius=3 cutoff=0 per/abs=0.03 link=1 displacement=6 dynamics=Brownian").

The analyses of the GEMs tracks were as described in (Delarue *et al*., 2018; Lemière *et al*., 2022), with methods to compute mean square displacement (MSD) using MATLAB (MATLAB_R2018, MathWorks). The effective diffusion D_eff_ was obtained by fitting the first 10 time points of the MSD curve (MSD_truncated_) to the canonical 2D diffusion law for Brownian motion: MSD_truncated_(τ)=4D_eff_ τ.

### Turgor pressure calculation

For the 3D Volume method, we fitted our data of protoplast volumes at various sorbitol concentrations to the Boyle Van’t Hoff equation for an ideal osmometer 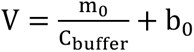 (Figure 2B, colored lines) where V is the volume of protoplast, C_%&’’()_ the concentration of the medium, m_;_ the apparent number of osmotically active particles, b_;_ the non-osmotic volume. m_;_, b_;_were parameters that were fitted to minimize the sum squares of the residues. Data from at least two experimental replicates were combined to generate these fits. Intact cells volume was the average value (±SD) of the median of the volume distribution of a population of cells from two replicates. The turgor pressure (Tables 1,2) was derived from estimating the sorbitol concentration in the Boyle Van’t Hoff plots that provided the equivalent median volume as in the intact cell population (Figure 2B). The estimated errors (Table 1,2) were computed by combining the variance of the fit from the protoplast data and the SD of median intact cell volume.

For cytoplasmic intensity method, we fitted the fluorescence intensity data in protoplasts at various sorbitol concentrations (Figure 3) to a simple linear regression close to the intensity values in the intact cells. The mean turgor pressure (Tables 1, 2) was estimated as the sorbitol concentration in the fitted equation that produced the equivalent mean fluorescence in the intact cell population. The estimated errors (Table 1, 2) were computed by combining the variance of the fit from the protoplast data with the variance of mean distributions of intact cell fluorescence intensity between replicates.

For the rheology method, we fit the measured *D_eff_* (effective diffusion coefficient) of 40-nm cytGEMs in protoplasts at various sorbitol concentrations to the Phillies equation *D*_*eff*_ = *D*_0_*e^-β(C+C_0_)^*, where *D*_0_ = 13.56 µm^2^/s is the diffusion of a 40 nm spherical particle in water calculated using the Stokes-Einstein equation, C_0_ the concentration of the medium, *C* the concentration of sorbitol (Figure 4) (see (Lemière *et al*., 2022)). The fit was derived from fitting the mean D_eff_ values from the combined data from all replicates. The mean turgor pressure (Tables 1, 2) was estimated as the sorbitol concentration in the Phillies fit that produced the equivalent mean *D_eff_* in the intact cell population. The estimated errors (Table 1, 2) were computed by combining the variance of the fit from the protoplast data with the standard error of the mean of intact cell *D_eff_* of all the tracks. For the Average values (Table1, 2, bottom rows), means of each of the three methods were reported, with the errors as the SD between these three values.

To compute the turgor pressure (P) from the concentration of sorbitol (C), we used the Van’t Hoff relation, P = CRT, where RT is the product of the gas constant, and the temperature is 303.15 K (30°C).

### Cell wall Young modulus and tension

To calculate the lateral cell wall modulus of fission yeasts we built upon the theoretical framework developed by (Atilgan *et al*., 2015). This framework assumes that the cell wall for *S. pombe and S. japonicus* is homogeneous and isotropic, which leads to a force balance equation that links the physical parameter of the cell to the elastic strain of the lateral cell wall:

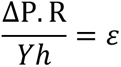

Where ΔP is the turgor pressure, and Y the Young modulus of the cell wall. We also used physical parameters for both fission yeast from (Davì *et al*., 2019), the measurements of the cell radius (R^Japonicus^ = 2 R^Pombe^), cell wall thickness (h^Japonicus^ = 2 h^Pombe^) and elastic strain (ε^Japonicus^ = ε^Pombe^ = 30%).

From the equation above and the differences in size between the two fission yeast species parameters it comes that:

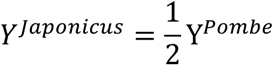

The cell wall tension (T) for a rod shape can be calculated following this equation:

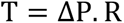

*S. japonicus* radius is twice *S. pombe* radius (R^Japonicus^ = 2R^Pombe^), but the turgor pressure bared by the former is only half of *S. pombe* turgor pressure 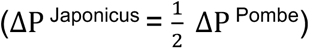. It leads to:

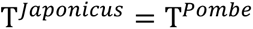

## Abbreviations used

S. pombe: Schizosaccharomyces pombe ;
S. japonicus: Schizosaccharomyces japonicus ;
S. cerevisiae: Saccharomyces cerevisiae;
D_eff_: effective diffusion,
cytGEMS: genetically-encoded multimeric cytoplasmic nanoparticles;
MSD: mean square displacement;
SD: standard deviation.

## Acknowledgments

We are grateful to members of Fred Chang’s lab as well as labs of Sophie Dumont, Orion Weiner, and David Drubin for helpful discussions. We thank Snezhka Oliferenko and Risa Mori for sharing materials and expertise for *S. japonicus* studies; Zhidong Tan for building the pAdh1sj::GFP::URA4sj plasmid used in this study, Jennifer Hill for providing concanavalin A, and Orna Cohen-Fix for providing *S. cerevisiae* strains and helpful advice. F.C. was supported by grants NIH 1R35GM141796 and NSF 2213582.

## Author Contributions

Conceptualization: J.L.; Methodology: J.L.; Software: J.L.; Validation: J.L.; Formal analysis: J.L.; Investigations: J.L.; Writing: J.L., F.C.; Review editing: J.L., F.C.; Visualization: J.L.; Supervision: J.L., F.C.; Project administration and funding acquisition: F.C.

## Notes

### Competing Interest Statement

The authors have declared no competing interest.

### Summary of Updates

Correction of typos, references added and addition of turgor pressure value on a fission yeast gpd1 mutant.

## Bibliography

Abenza, JF, Couturier, E, Dodgson, J, Dickmann, J, Chessel, A, Dumais, J, and Carazo Salas, RE (2015). Wall mechanics and exocytosis define the shape of growth domains in fission yeast. Nat Commun.

Aghamohammadzadeh, S, and Ayscough, KR (2009). Differential requirements for actin during yeast and mammalian endocytosis. Nat Cell Biol 11, 1039–1042.

Aiba, H, Yamada, H, Ohmiya, R, and Mizuno, T (1995). The osmo-inducible gpdl+ gene is a target of the signaling pathway involving Wis1 MAP-kinase kinase in fission yeast. FEBS Lett 376, 199– 201.

Altenburg, T, Goldenbogen, B, Uhlendorf, J, and Klipp, E (2019). Osmolyte homeostasis controls single-cell growth rate and maximum cell size of Saccharomyces cerevisiae. Npj Syst Biol Appl 5, 34.

Aoki, K, and Niki, H (2017). Transformation of Schizosaccharomyces japonicus. Cold Spring Harb Protoc 2017, 996–998.

Arnold, WN, and Lacy, JS (1977). Permeability of the cell envelope and osmotic behavior in Saccharomyces cerevisiae. J Bacteriol 131, 564–571.

Atilgan, E, Magidson, V, Khodjakov, A, and Chang, F (2015). Morphogenesis of the Fission Yeast Cell through Cell Wall Expansion. Curr Biol 25, 2150–2157.

Bähler, J, Wu, J, Longtine, MS, Shah, NG, Mckenzie III, A, Steever, AB, Wach, A, Philippsen, P, and Pringle, JR (1998). Heterologous modules for efficient and versatile PCR-based gene targeting in Schizosaccharomyces pombe. Yeast 14, 943–951.

Basu, R, Munteanu, EL, and Chang, F (2013). Role of turgor pressure in endocytosis in fission yeast. Mol Biol Cell 25, 679–687.

Beauzamy, L, Derr, J, and Boudaoud, A (2015a). Quantifying hydrostatic pressure in plant cells by using indentation with an atomic force microscope. Biophys J 108, 2448–2456.

Beauzamy, L, Louveaux, M, Hamant, O, and Boudaoud, A (2015b). Mechanically, the shoot apical meristem of arabidopsis behaves like a shell inflated by a pressure of about 1 MPa. Front Plant Sci 6, 1–10.

Beauzamy, L, Nakayama, N, and Boudaoud, A (2014). Flowers under pressure: Ins and outs of turgor regulation in development. Ann Bot.

Boulbitch, AA (1998). Deflection of a cell membrane under application of a local force. Phys Rev E - Stat Physics, Plasmas, Fluids, Relat Interdiscip Top 57, 2123–2128.

Brachmann, CB, Davies, A, Cost, GJ, Caputo, E, Li, J, Hieter, P, and Boeke, JD (1998). Designer deletion strains derived from Saccharomyces cerevisiae S288C: a useful set of strains and plasmids for PCR-mediated gene disruption and other applications. Yeast 14, 115–132.

Chang, F, and Huang, KC (2014). How and why cells grow as rods. BMC Biol 12, 54.

Cherry, JM et al. (2012). Saccharomyces Genome Database: The genomics resource of budding yeast. Nucleic Acids Res 40, 700–705.

Cohen, R, and Engelberg, D (2007). Commonly used Saccharomyces cerevisiae strains (e.g. BY4741, W303) are growth sensitive on synthetic complete medium due to poor leucine uptake. FEMS Microbiol Lett 273, 239–243.

Davì, V, Chevalier, L, Guo, H, Tanimoto, H, Barrett, K, Couturier, E, Boudaoud, A, and Minc, N (2019). Systematic mapping of cell wall mechanics in the regulation of cell morphogenesis. Proc Natl Acad Sci U S A 116, 13833–13838.

Davì, V, Tanimoto, H, Ershov, D, Haupt, A, De Belly, H, Le Borgne, R, Couturier, E, Boudaoud, A, and Minc, N (2018). Mechanosensation Dynamically Coordinates Polar Growth and Cell Wall Assembly to Promote Cell Survival. Dev Cell 45, 170–182.e7.

Delarue, M et al. (2018). mTORC1 Controls Phase Separation and the Biophysical Properties of the Cytoplasm by Tuning Crowding. Cell 174, 338–349.e20.

Dmitrieff, S, and Nédélec, F (2015). Membrane Mechanics of Endocytosis in Cells with Turgor. PLoS Comput Biol 11.

Edelstein, A, Amodaj, N, Hoover, K, Vale, R, and Stuurman, N (2010). Computer control of microscopes using manager. Curr Protoc Mol Biol 92.

Edelstein, AD, Tsuchida, MA, Amodaj, N, Pinkard, H, Vale, RD, and Stuurman, N (2014). Advanced methods of microscope control using μManager software. J Biol Methods 1, e10.

Emmett, RW, and Parbery, DG (1975). Appressoria. Annu Rev Phytopathol 13, 147–165.

Flor-Parra, I, Zhurinsky, J, Bernal, M, Gallardo, P, and Daga, RR (2014). A Lallzyme MMX-based rapid method for fission yeast protoplast preparation. Yeast 31, 61–66.

Goldenbogen, B, Giese, W, Hemmen, M, Uhlendorf, J, Herrmann, A, and Klipp, E (2016). Dynamics of cell wall elasticity pattern shapes the cell during yeast mating morphogenesis. Open Biol 6.

Huh, WK, Falvo, J V., Gerke, LC, Carroll, AS, Howson, RW, Weissman, JS, and O’Shea, EK (2003). Global analysis of protein localization in budding yeast. Nature 425, 686–691.

Jiang, H, and Sun, SX (2010). Morphology, growth, and size limit of bacterial cells. Phys Rev Lett 105, 1–4.

Jones, TM, Marks, PC, Cowan, JM, Kainth, DK, and Petrie, RJ (2021). Cytoplasmic pressure maintains epithelial integrity and inhibits cell motility. Phys Biol 18.

Kashiwazaki, J, Yamasaki, Y, Itadani, A, Teraguchi, E, Maeda, Y, Shimoda, C, and Nakamura, T (2011). Endocytosis is essential for dynamic translocation of a syntaxin 1 orthologue during fission yeast meiosis. Mol Biol Cell 22, 3658–3670.

Lang, F, Busch, GL, Ritter, M, Völkl, H, Waldegger, S, Gulbins, E, and Häussinger, D (1998). Functional significance of cell volume regulatory mechanisms. Physiol Rev 78, 247–306.

Lemière, J, Real-Calderon, P, Holt, LJ, Fai, TG, and Chang, F (2022). Control of nuclear size by osmotic forces in Schizosaccharomyces pombe. Elife 11.

Lemière, J, Ren, Y, and Berro, J (2021). Rapid adaptation of endocytosis, exocytosis and eisosomes after an acute increase in membrane tension in yeast cells. Elife 10, 1–29.

Li, Y, He, L, Gonzalez, NAP, Graham, J, Wolgemuth, C, Wirtz, D, and Sun, SX (2017). Going with the Flow: Water Flux and Cell Shape during Cytokinesis. Biophys J 113, 2487–2495.

Longtine, MS, McKenzie, A, Demarini, DJ, Shah, NG, Wach, A, Brachat, A, Philippsen, P, and Pringle, JR (1998). Additional modules for versatile and economical PCR-based gene deletion and modification in Saccharomyces cerevisiae. Yeast 14, 953–961.

Machado, S, Mercier, V, and Chiaruttini, N (2019). LimeSeg: a coarse-grained lipid membrane simulation for 3D image segmentation. BMC Bioinformatics 20, 2.

Martinez de Marañon, I, Marechal, P-A, and Gervais, P (1996). Passive Response of Saccharomyces cerevisiaeto Osmotic Shifts: Cell Volume Variations Depending on the Physiological State. Biochem Biophys Res Commun 227, 519–523.

Matheson, K, Parsons, L, and Gammie, A (2017). Whole-genome sequence and variant analysis of W303, a widely-used strain of Saccharomyces cerevisiae. G3 Genes, Genomes, Genet 7, 2219–2226.

Meikle, AJ, Reed, RH, and Gadd, GM (2009). Osmotic Adjustment and the Accumulation of Organic Solutes in Whole Cells and Protoplasts of Saccharomyces cerevisiae. Microbiology 134, 3049–3060.

Milo, R, and Phillips, R (2015). Cell Biology by the Numbers, Garland Science.

Minc, N, Boudaoud, A, and Chang, F (2009). Mechanical Forces of Fission Yeast Growth. Curr Biol 19, 1096–1101.

Minton, AP (1981). Excluded volume as a determinant of macromolecular structure and reactivity. Biopolymers 20, 2093–2120.

Mishra, R, Minc, N, and Peter, M (2022). Cells under pressure: how yeast cells respond to mechanical forces. Trends Microbiol 30, 495–510.

Molines, AT et al. (2022). Physical properties of the cytoplasm modulate the rates of microtubule polymerization and depolymerization. Dev Cell 57, 466–479.e6.

Mortimer, RK, and Johnston, JR (1986). Genealogy of principal strains of the yeast genetic stock center. Genetics 113, 35–43.

Nickaeen, M, Berro, J, Pollard, TD, and Slepchenko, BM (2022). A model of actin-driven endocytosis explains differences of endocytic motility in budding and fission yeast. Mol Biol Cell 33, ar16.

Nobel, PS (1969). The Boyle-Van’t Hoff relation. J Theor Biol.

Petrezselyova, S, Zahradka, J, and Sychrova, H (2010). Saccharomyces cerevisiae BY4741 and W303-1A laboratory strains differ in salt tolerance. Fungal Biol 114, 144–150.

Phillies, GDJ (1988). Quantitative prediction of .alpha. in the scaling law for self-diffusion. Macromolecules 21, 3101–3106.

Proctor, SA, Minc, N, Boudaoud, A, and Chang, F (2012). Contributions of turgor pressure, the contractile ring, and septum assembly to forces in cytokinesis in fission yeast. Curr Biol 22, 1601–1608.

Ralser, M, Kuhl, H, Ralser, M, Werber, M, Lehrach, H, Breitenbach, M, and Timmermann, B (2012). The Saccharomyces cerevisiae W303-K6001 cross-platform genome sequence: Insights into ancestry and physiology of a laboratory mutt. Open Biol 2.

Rogowska-Wrzesinska, A, Larsen, PM, Blomberg, A, Görg, A, Roepstorff, P, Norbeck, J, and Fey, SJ (2001). Comparison of the proteomes of three yeast wild type strains: CEN.PK2, FY1679 and W303. Comp Funct Genomics 2, 207–225.

Rothstein, RJ, and Sherman, F (1980a). Dependence on mating type for the overproduction of iso-2-cytochrome c in the yeast mutant CYC7-H2. Genetics 94, 891–898.

Rothstein, RJ, and Sherman, F (1980b). Genes affecting the expression of cytochrome c in yeast: genetic mapping and genetic interactions. Genetics 94, 871–889.

Routier-Kierzkowska, A-L, Weber, A, Kochova, P, Felekis, D, Nelson, BJ, Kuhlemeier, C, and Smith, RS (2012). Cellular Force Microscopy for in Vivo Measurements of Plant Tissue Mechanics. Plant Physiol 158, 1514–1522.

Sbalzarini, IF, and Koumoutsakos, P (2005). Feature point tracking and trajectory analysis for video imaging in cell biology. J Struct Biol 151, 182–195.

Schneider, CA, Rasband, WS, and Eliceiri, KW (2012). NIH Image to ImageJ: 25 years of image analysis. Nat Methods 9, 671–675.

Shabala, L, McMeekin, T, and Shabala, S (2009). Osmotic adjustment and requirement for sodium in marine protist thraustochytrid. Environ Microbiol 11, 1835–1843.

Thilini Chethana, KW, et al. (2021). Diversity and function of appressoria. Pathogens 10, 3–6.

Tomos, AD, and Leigh, RA (1999). The pressure probe: A versatile tool in plant cell physiology. Annu Rev Plant Biol 50, 447–472.

Tsugawa, S et al. (2022). Elastic shell theory for plant cell wall stiffness reveals contributions of cell wall elasticity and turgor pressure in AFM measurement. Sci Rep 12, 13044.

Vian, A, Pochitaloff, M, Yen, S-T, Kim, S, Pollock, J, Liu, Y, Sletten, EM, and Campas, O (2022). In situ quantification of osmotic pressure within living embryonic tissues. BioRxiv, 2022.12.04.519060.

Wagner, E, and Glotzer, M (2016). Local RhoA activation induces cytokinetic furrows independent of spindle position and cell cycle stage. J Cell Biol 213, 641–649.

Wang, X, Li, L, Shao, Y, Wei, J, Song, R, Zheng, S, Li, Y, and Song, F (2021). Effects of the Laplace pressure on the cells during cytokinesis. IScience 24, 102945.

Wieland, J, Nitsche, AM, Strayle, J, Steiner, H, and Rudolph, HK (1995). The PMR2 gene cluster encodes functionally distinct isoforms of a putative Na+ pump in the yeast plasma membrane. EMBO J 14, 3870–3882.

Yang, NJ, and Hinner, MJ (2015). Getting Across the Cell Membrane: An Overview for Small Molecules, Peptides, and Proteins. In: Site-Specific Protein Labeling: Methods and Protocols, 29–53.

Zadrag-Tecza, R, Kwolek-Mirek, M, Bartosz, G, and Bilinski, T (2009). Cell volume as a factor limiting the replicative lifespan of the yeast Saccharomyces cerevisiae. Biogerontology 10, 481– 488.

